# Generative modeling of single-cell population time series for inferring cell differentiation landscapes

**DOI:** 10.1101/2020.08.26.269332

**Authors:** Grace H.T. Yeo, Sachit D. Saksena, David K. Gifford

## Abstract

Existing computational methods that use single-cell RNA-sequencing for cell fate prediction either summarize observations of cell states and their couplings without modeling the underlying differentiation process, or are limited in their capacity to model complex differentiation landscapes. Thus, contemporary methods cannot predict how cells evolve stochastically and in physical time from an arbitrary starting expression state, nor can they model the cell fate consequences of gene expression perturbations. We introduce PRESCIENT (Potential eneRgy undErlying Single Cell gradIENTs), a generative modeling framework that learns an underlying differentiation landscape from single-cell time-series gene expression data. Our generative model framework provides insight into the process of differentiation and can simulate differentiation trajectories for arbitrary gene expression progenitor states. We validate our method on a recently published experimental lineage tracing dataset that provides observed trajectories. We show that this model is able to predict the fate biases of progenitor cells in neutrophil/macrophage lineages when accounting for cell proliferation, improving upon the best-performing existing method. We also show how a model can predict trajectories for cells not found in the model’s training set, including cells in which genes or sets of genes have been perturbed. PRESCIENT is able to accommodate complex perturbations of multiple genes, at different time points and from different starting cell populations. PRESCIENT models are able to recover the expected effects of known modulators of cell fate in hematopoiesis and pancreatic β cell differentiation.

## Introduction

Waddington’s epigenetic landscape posits that the state of differentiating cells can be modeled by balls rolling down a hilly landscape toward valleys corresponding to cell fates. In this view, topography of the differentiation landscape corresponds to biological ‘potential energy’, with cells traveling from regions of high energy toward regions of low energy (Ferrell, 2012). Mapping this landscape is essential to improving our understanding of how cells are driven to transient and terminal states *in vivo* and to enable precise manipulation of cell fates *in vitro* (Cohen and Melton, 2011).

Single-cell RNA-sequencing (scRNA-seq) technology has enabled the study of developmental landscapes by the observation of gene expression in single cells sampled at multiple stages of differentiation. However, these studies provide snapshots of heterogeneous cell states of a given differentiation process and typically do not directly observe lineage relationships between cells. Furthermore, because cells are destroyed in order to collect transcriptomic information, the ancestors and children of any observed cell are not measured. Recently, experimental lineage tracing methods that couple various barcoding strategies with scRNA-seq have been described that identify relationships between cells (Biddy et al., 2018; Wagner and Klein, 2020; Weinreb et al., 2020). However, these methods are still in relative infancy and have not yet been widely adopted.

Computational methods have also been developed to infer cell lineages from scRNA-seq data in the absence of experimental tracing (Figure 1a). However, these methods typically summarize observations of cell states and couplings emergent of the underlying process and have limited to no capacity for modeling differentiation as a stochastic process in physical time (Saelens et al., 2019; Tanay and Regev, 2017; Trapnell et al., 2012). The predominant approach is pseudo-temporal inference, which orders cells along an arbitrary one-dimensional measurement representing ‘differentiation time’, and hence cannot model differentiation dynamics with respect to real, physical time. Other methods have also emerged for the specific task of cell fate prediction. For example, Waddington-OT predicts long-range cell-cell probabilistic couplings by reframing the task of inferring cell relationships between population snapshots as an unbalanced optimal transport problem (Schiebinger et al., 2019). Another method, Fate-ID iteratively builds ensembled cell-type classifiers from labeled terminal cell states (Herman et al., 2018). However, these methods only summarize observations of cell states and couplings emergent of the underlying differentiation process.

**Figure 1.**
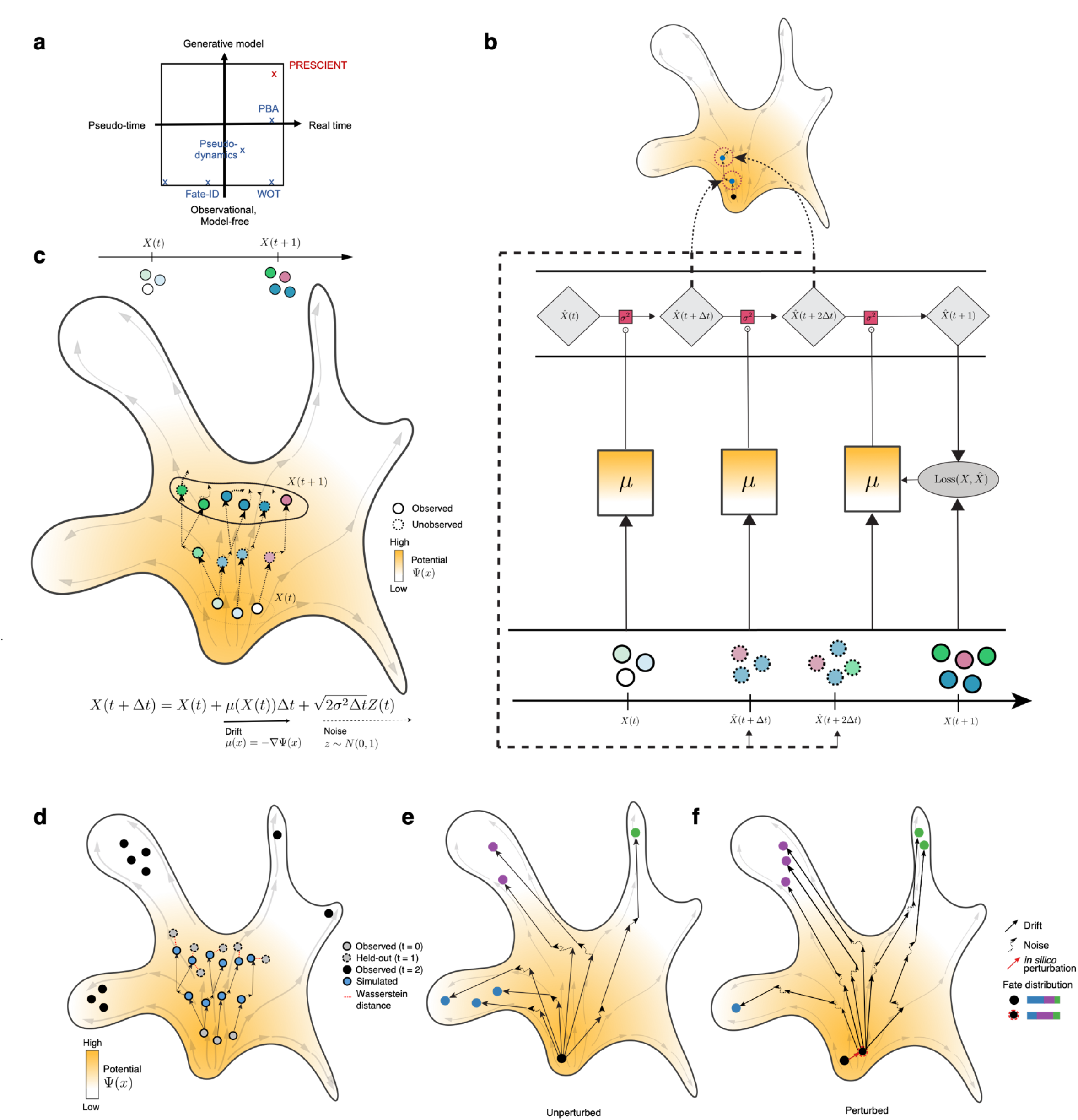
A generative model of cellular differentiation. (a) Existing single-cell models of development can be classified as operating in pseudo-time or real time (x-axis), and the extent to which the method models the underlying differentiation process (y-axis). PRESCIENT is shown in red (b) Observations of population-level time-series data are used in a generative framework that models the underlying dynamic process in physical time. Evolution of a cell’s state is governed by a drift term and a noise term. The drift, depicted by solid arrows, is defined as the negative gradient of the potential function, depicted by the color gradient in the background. Dashed lines correspond to noise. The model is fit using observations of population-level time-series data, depicted as solid circles. Simulations of cell states are depicted as dashed circles. (c) Cartoon depicting model fitting process. The neural network parameterizing the underlying drift function *μ* takes as input the PCA projections of gene expression data at observed time points (again depicted as solid lines). The stochastic process is then simulated via first-order time discretization to produce a population at the next time step, and so on. This proceeds until the next observed time point, at which the loss between the simulated and predicted population is minimized. The model was validated using two tasks (d) held-out recovery, where the model was asked to predict the marginal distribution of a held-out time point, and (e) fate prediction, where the model was asked to predict the fate distribution outcome of a given progenitor cell. Fate prediction can be applied to cells observed in the dataset (left) or cell states in which some perturbation has been applied *in silico* (right). As shown, the perturbation results in a shift of fate distribution outcomes.

Recently, a small number of methods have described approaches to modeling differentiation as a process, but they have been limited either in how the model is solved, or in modeling capacity. For example, Population Balance Analysis (PBA) solves a reaction-diffusion partial differential equation describing differentiation but is forced to use a non-parametric solution due to computational constraints (Weinreb et al., 2018). Similarly, pseudo-dynamics models a diffusion process but only in a one-dimensional cell state (Fischer et al., 2019). Neither have shown to be able to simulate differentiation trajectories starting from an unseen cell state. However, a model that is able to generalize to data points outside of the training set is required to be able to make predictions for the consequences of *in silico* perturbations that can result in a cell state that has not yet been observed.

We build upon a modeling framework by Hashimoto et al. in which cellular differentiation is modeled as a diffusion process over a potential landscape parameterized by a neural network (Hashimoto et al., 2016). Neural networks are powerful black-box function approximators that allow arbitrary and extremely complex parameterizations of the landscape. Since this modeling framework is fully generative, it is naturally able to sample trajectories for data points not present in the training set. However, the original work was limited to considering relatively simple models and small scRNA-seq datasets with up to 10 genes. It also did not evaluate trajectories sampled from the model and did not take into account cell proliferation, which is well-known to be an important factor in cellular differentiation (Ruijtenberg and van den Heuvel, 2016; Schiebinger et al., 2019).

We introduce PRESCIENT (**P**otential ene**R**gy und**E**rlying **S**ingle **C**ell grad**IENT**s), a generative modeling framework that is fit using longitudinal scRNA-seq datasets that include irregularly sampled time-series with large numbers of cells and high-dimensional measurements. We validate our model on a newly published lineage tracing dataset. In particular, we evaluate our model on its ability to recover held-out time points, and to predict cell fate biases established by experimental lineage tracing. We build upon previous work in three ways. First, we study different parameterizations of underlying potential functions, and show that more complex models can improve the performance of recovering a held-out time point. Second, we extend the modeling framework to incorporate cell proliferation, and show that taking into account proliferation is crucial for fate prediction. Third, we demonstrate how PRESCIENT can be used to model perturbations of multiple genes, at different time points, and from different starting states. We are able to recover expected changes in final cell fate distributions when interrogating our models using perturbations of known regulators of cell fate in hematopoiesis and pancreatic β cell differentiation. This enables large unbiased *in silico* perturbation experiments, both individual and combinatorial, to aid the design of *in vitro* genetic perturbational screens.

## Results

### Learning a generative model of cellular differentiation from high-dimensional scRNA-seq data

PRESCIENT models cellular differentiation as a diffusion process over a gene expression landscape parameterized by a potential function that we wish to identify given only time-series population snapshots of single-cell RNA expression. In this diffusion process, evolution of a cell’s state at a given time is governed by a drift term, corresponding to the force acting on that cell given its current state, and a noise term, corresponding to stochasticity. In particular, we consider the case where the drift term is defined to be the negative gradient of the potential function, such that the potential induces a force that naturally drives cells toward regions of low potential (Figure 1b; Methods). This stochastic process can then be simulated via first-order time discretization to sample trajectories for a given cell. The potential function is fit by minimizing a regularized Wasserstein loss between empirical and predicted populations at the observed time points (Figure 1c; Methods) (Hashimoto et al., 2016).

To enable the modeling framework to operate on large scRNA-seq datasets, we fit models on PCA projections of the scaled gene expression data rather than on the gene expression itself. Euclidean distance in PCA space has been successfully used in down-stream scRNA-seq analysis methods such as clustering and cross-dataset integration and thus use it as a reasonable distance metric for scRNA-seq data. We also take advantage of modern deep learning methods to study alternative parameterizations of the underlying potential function. We use automatic differentiation of a neural network potential function to calculate potential function gradients to determine the associated drift function of the model.

Finally, we extend the framework to take into account cell proliferation by weighting each cell in the source population according to its expected number of descendants in the objective function. This is in contrast to Waddington-OT which uses estimates of cell proliferation to instead formulate an unbalanced optimal transport problem. To assess the importance of incorporating cell proliferation, we study models assuming *a priori* knowledge of cell proliferation, which can be directly estimated from the data by computing the number of descendants for each starting cell given lineage tracing data. However, since we are largely concerned with model estimation in the absence of lineage tracing, we also study models where cell proliferation is estimated from gene expression.

We first validate our model on a recently published lineage tracing dataset by Weinreb et al. which used DNA barcodes to track clonal trajectories during mouse hematopoiesis (Weinreb et al., 2020). We evaluate our model on two tasks: recovery of a population at a held-out time point, and cell fate prediction (Figure 1d-e).

### Generative modeling recovers population state at a held-out time point

Deep learning enables the modeling of complex potential functions with correspondingly large parameter sets. We hypothesized that more expressive parameterizations of the potential function might improve modeling capacity of the diffusion process. We hence evaluated potential functions of differing complexity on the task of predicting the marginal distribution of gene expression at a held-out time point.

We fit models with four different architectures (fully connected neural network models with 1 layer of 1000 units, 1 layer of 4000 units, 2 layers of 200 units, or 2 layers of 400 units, all using softplus as the activation function), and using three different regularization strengths (tau = 1e-3, 1e-6, 1e-9). These architectures were chosen such that 2-layer models could be compared with a 1-layer model with roughly the same number of parameters. Models were fit via pre-training on day 6, and then trained on days 2 and 6. In particular, we used only the subset of data for which lineage tracing data is available to enable comparison of models with and without ground truth cell proliferation rates. For evaluation, we computed the Wasserstein distance between the simulated cell population and the empirically observed cell population at days 4 (training) and 6 (testing) (Figure 2a). The testing distance was evaluated at the epoch with the lowest training distance (Figure S1a-b).

**Figure 2.**
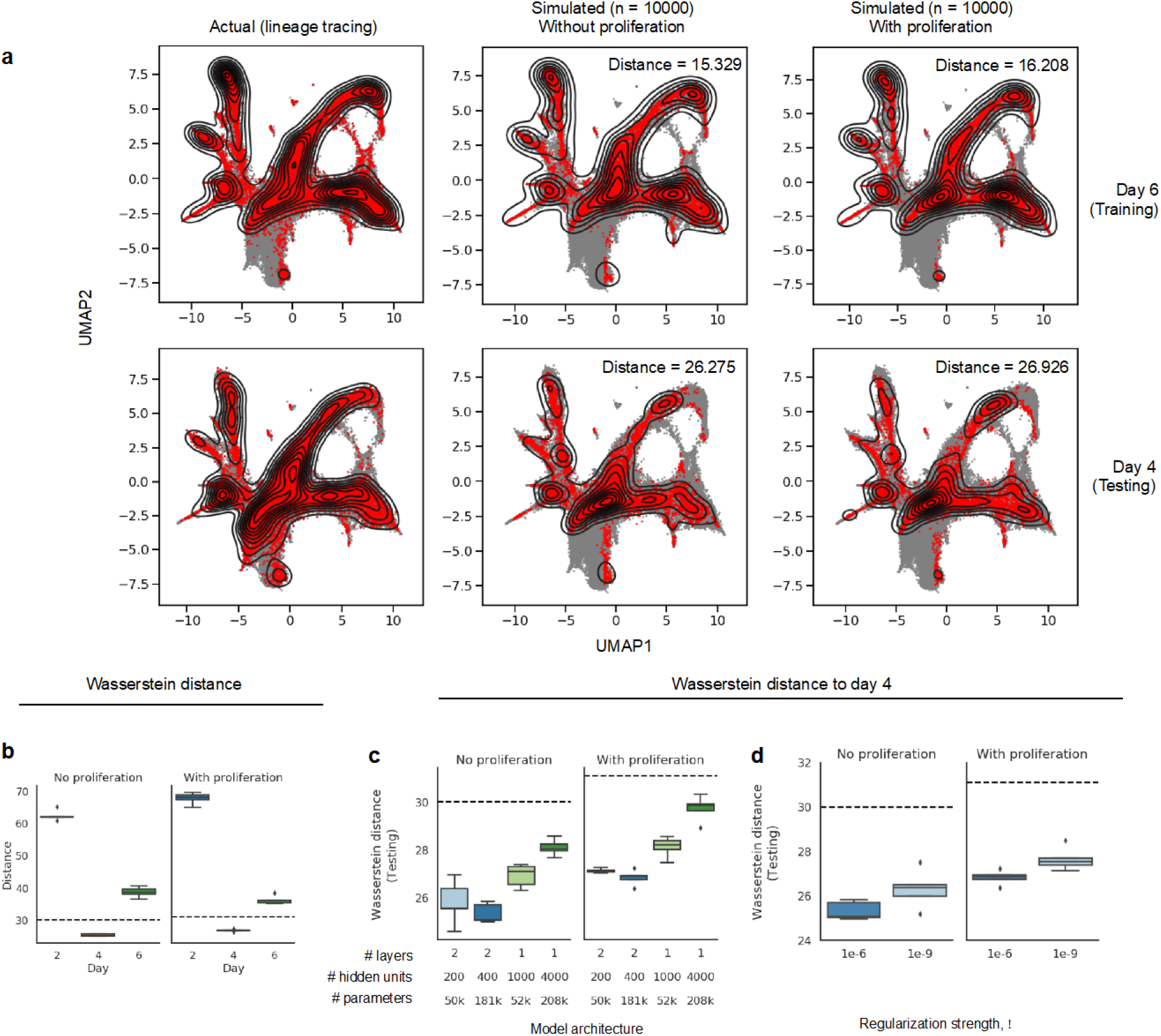
Generative modeling framework recovers population at held-out time point. (a-b) Results for 2-layer models with 400 units and a regularization strength of tau = 1e-6, either with or without cell proliferation (a) Actual and simulated populations on days 6 (training, top), and 4 (testing, below). Distance reported is the Wasserstein distance of the simulated population to the actual population at the given time point. (b) Wasserstein distance of simulated population at day 4 to actual populations at each time point (b-d) Dashed line indicates distance of linear interpolated population with respect to actual population at day 4 (c-d) Testing performance for models of different complexity (c) and with different regularization strengths (d) for 5 seeds.

As baselines, we also computed the distance of the simulated population at day 4 to the actual populations at day 2 and 6. We next computed the distance of the actual population at day 4 to a linearly interpolated population as implemented by Waddington-OT (indicated on plots by a dashed line) (Figure 2b). Note that we should expect linear interpolation to perform well when populations are sampled at dense time-points. Overall, all models out-performed these baselines except when the regularization strength was too high (i.e. except when *τ* = 1e-3), (Figure 2b-d, S1d).

For both models trained with or without cell proliferation, we observed that although 1-layer models may perform as well as 2-layer models on training, 2-layer models achieve lower testing distance even when the number of parameters in the 1-layer models was greater than or equivalent to in the 2-layer models (Figure 2c, S1b-c). This could suggest that the 2-layer models are either better able to approximate the potential function, or are easier to fit. We also observed that model training and testing distance was best with a moderate regularization strength of *τ* =1e-6 across model architectures (Figure 2d, Figure S1c-d). In general, models accounting for cell proliferation performed slightly worse at predicting the held-out time point than models that did not account for cell proliferation.

We reasoned that hyper-parameters selected on this task would be transferable to models for fate prediction because good recovery of the held-out time point should imply that the model had been able to find a good approximation of the underlying potential function. We hence use 2-layer models with 400 units and a regularization strength of 1e-6 for subsequent tasks.

### Fate outcomes generated by PRESCIENT align with experimental lineage tracing when taking into account cell proliferation

We next sought to investigate if the PRESCIENT modeling framework improves performance on predicting cell fate. Broadly, cell fate prediction is the task of predicting the final distribution of cell states that a cell reaches given its state at an earlier time point. For this dataset, we are concerned in particular with the task of predicting the relative distributions of neutrophil and monocyte cells at day 6 of a given cell lineage given a cell from that lineage at day 2. In particular, we compared model performance with and without incorporating cell proliferation.

We consider the clonal fate bias metric described by Weinreb et al., defined as the number of neutrophils divided by the total number of neutrophils and monocytes. For each cell in the test set, we simulate 2000 trajectories until the final time point. For each of these trajectories, we classify the cell at the final time point as neutrophil, monocyte or other using an approximate nearest neighbor (ANN) classifier that had been trained using cell-type labels provided by Weinreb et al. (Methods). We then evaluate the clonal fate bias on the test set, consisting of 335 cells on Day 2 (Figure 3a,c). To reduce noise, we ensemble predictions by taking the mean of the clonal fate bias metric over the last 5 evaluated epochs (Figure S1e). We measure performance as the Pearson correlation with respect to the actual clonal fate bias given by the lineage tracing data. Since we observed that the clonal fate bias as measured in the lineage tracing data was strongly bimodal (Figure S1g), we also considered a second metric, which we define to be the AUROC of classifying a given cell as having a clonal fate bias of > 0.5.

**Figure 3.**
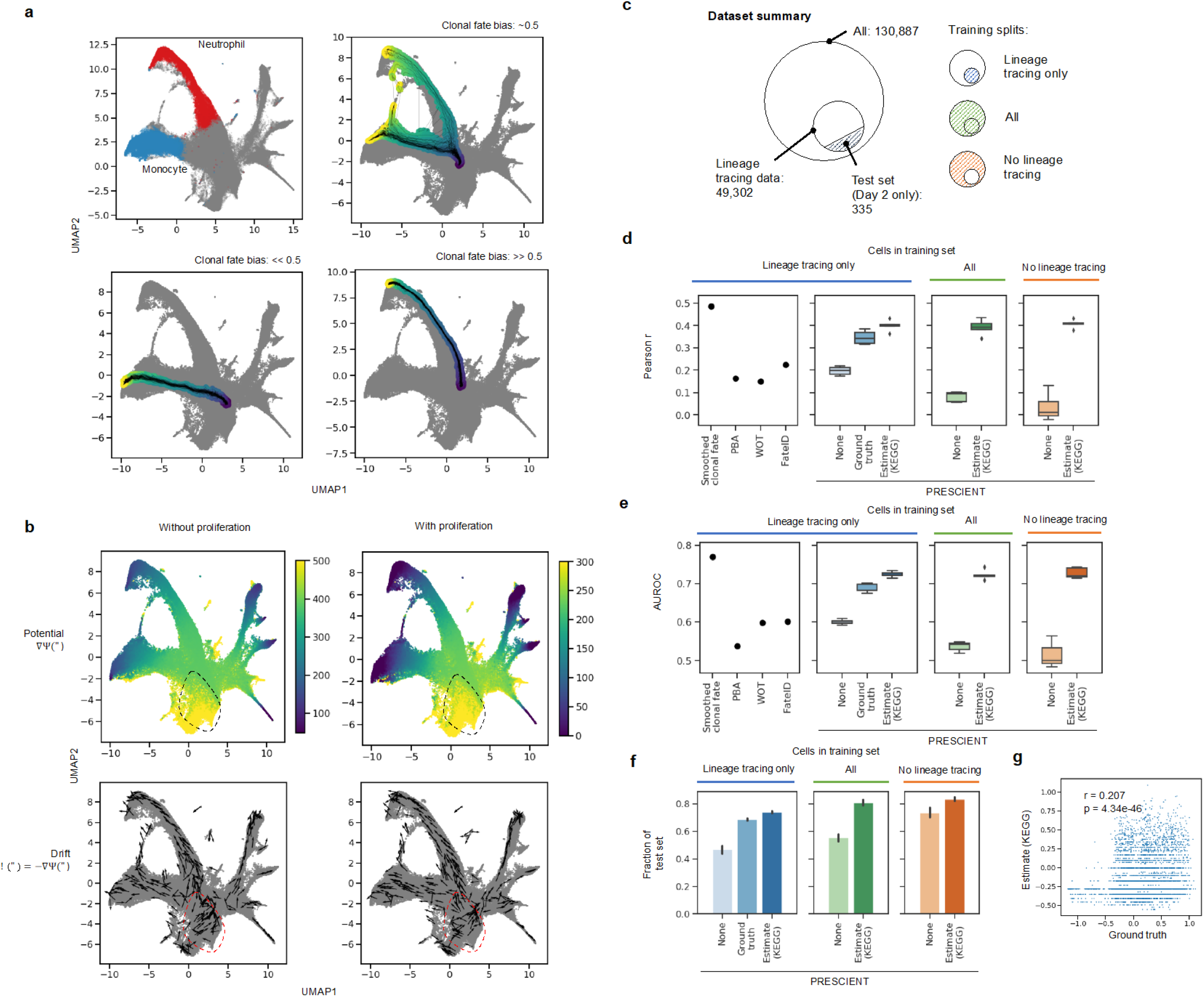
Incorporating cell proliferation greatly improves model performance on predicting cell fate. (a) Examples of trajectories with starting cells assigned different clonal fate biases. The colorscale indicates time with darker (purple) corresponding to t = 2 and lighter (yellow) corresponding to t = 6. Clonal fate bias is computed with respect to monocyte/neutrophil populations at the final time point (b) Visualizations of underlying potential and drift functions learned by models with and without cell proliferation. Drift is visualized for a random sample of cells. Dotted circle indicates qualitative differences in potential landscape. (c) Summary of training/test splits of lineage tracing dataset. Training splits either include only cells with lineage tracing data, all cells, or cells without lineage tracing data. (d-f) Performance of other methods (far left, Smoothed fate probabilities given by held-out clonal data, PBA: population balance analysis, WOT: Waddington-OT; evaluated using predictions provided by Weinreb et al.) in comparison to PRESCIENT (left-right) on predicting clonal fate bias for the same test set given different training sets. Training sets either consisted of only cells for which lineage tracing data was available (left), all cells (middle), or cells without lineage tracing data (right). Performance metrics evaluated include (d) Pearson r (e) AUROC, and (f) fraction of test set in which at least 1 simulated cell at the final time point is either classified as a neutrophil or a monocyte (g) Correlation of actual and estimated proliferation rates.

We first fit a PRESCIENT model on the subset of data for which lineage tracing data is available (Figure 3c). Figure 3b depicts the learned potential and drift functions for models trained with and without cell proliferation. Qualitatively, we observe that incorporating cell proliferation changes the potential landscape near the earliest time point. When the model does not consider cell proliferation, its performance is similar to existing fate prediction methods like Waddington-OT with provided ground truth cell proliferation (r = 0.150, AUROC = 0.599) and FateID (r = 0.225, AUROC = 0.602), achieving r = 0.196 +/- 0.020, AUROC = 0.601 +/- 0.006 over 5 seeds. Incorporating ground truth cell proliferation rates into the PRESCIENT modeling framework greatly improves performance with mean r = 0.347 +/- 0.029, AUROC = 0.692 +/- 0.012, more closely approaching the upper-bound performance estimated using held-out clonal data of r = 0.487, AUROC = 0.771 (Figure 3d-f).

We have shown that accounting for cell proliferation in the modeling framework improves the performance of the fate prediction task. However, these models were fit with ground truth cell proliferation rates derived from the lineage tracing data, which is not only generally unavailable for most scRNA-seq time-series datasets, but even missing for 62.3% of this dataset (Figure 3c). We next looked at whether cell proliferation rates derived from gene expression could achieve similar performance. To this end, we modified an approach originally described by Schiebinger et al. to use KEGG annotations of cell cycle and apoptosis genes which were also highly variable in the dataset to estimate the number of descendants a cell is expected to have. These estimates correlated weakly but significantly (r = 0.207, p << 1e-45) with the ground-truth rates calculated from the lineage tracing data (Figure 3g, S1d). We compared models fit to the same set of cells using either our gene-expression-derived or ground-truth proliferation estimates. We found that models incorporating gene-expression-derived estimates achieved r = 0.399 +/- 0.025 and AUROC = 0.725 +/- 0.008, a slight improvement over ground-truth proliferation -based models (Figure 3d-e).

### Generative models can predict cell states not observed during training

We next hypothesized that PRESCIENT should be able to predict the fate of cells not observed during training. We expect that a PRESCIENT model has learned a good approximation of the underlying potential function and hence should be able to generalize to data points that were not observed during training. To test our hypothesis, we used our proliferation estimates to fit models to all cells with and without lineage tracing data.

We found that models incorporating estimated proliferation rates vastly outperformed models that did not in predicting the future expression of the previously described test set. We also found that model performance was similar when the cells in the test set were included in the training dataset (r = 0.391 +/- 0.035, AUROC = 0.723 +/- 0.013), and when they were not (r = 0.407 +/- 0.019, AUROC = 0.727 +/- 0.014) (Figure 3c-e).

Finally, although performance as measured by correlation and AUROC is similar to when models were fit only on cells with lineage tracing data, the fraction of test set cells for which the model predicted at least one cell entering a neutrophil or monocyte fate increased slightly from 0.74 +/- 0.01 to 0.80 +/- 0.03 (all cells) and 0.83 +/- 0.02 (only cells without lineage tracing data) (Figure 3c,f), suggesting that the models did benefit from observing more data.

### Computational gene expression perturbations produce expected changes in cell fates

The ability to simulate trajectories for out-of-sample cells allows the model to make predictions of the fate distribution outcome of cells with perturbed gene expression profiles. We demonstrate this ability on two model systems drawn from published studies with time-series scRNA-seq measurements of differentiation. For these experiments, we sampled a starting cell population of 200 cells from the earliest time point. We use this starting cell population to sample model predicted trajectories for an unperturbed population. We then introduce perturbations to the same starting cell population by simulating different levels of overexpression or knock-down/out of either individual genes or small subsets of genes that have been reported in the literature to modulate cell fate outcome in differentiation trajectories. We do this by setting the scaled gene-expression of the target gene to a z-score of less than zero for knockdowns and greater than zero for overexpression. These perturbations can involve multiple genes, or be introduced at different time points, or in different starting populations. The resulting gene expression profile is then transformed into PCA space to sample trajectories for the perturbed population. At each time step, we classify cell types using an ANN classifier trained on cell type labels derived from the original study. Finally, to test if a perturbation has had a significant effect on cell fate bias, we conduct t-tests comparing the proportions of a given cell fate at the final time point in the perturbed and unperturbed populations over 10 random initializations (Figure 4a-b, 5b).

**Figure 4.**
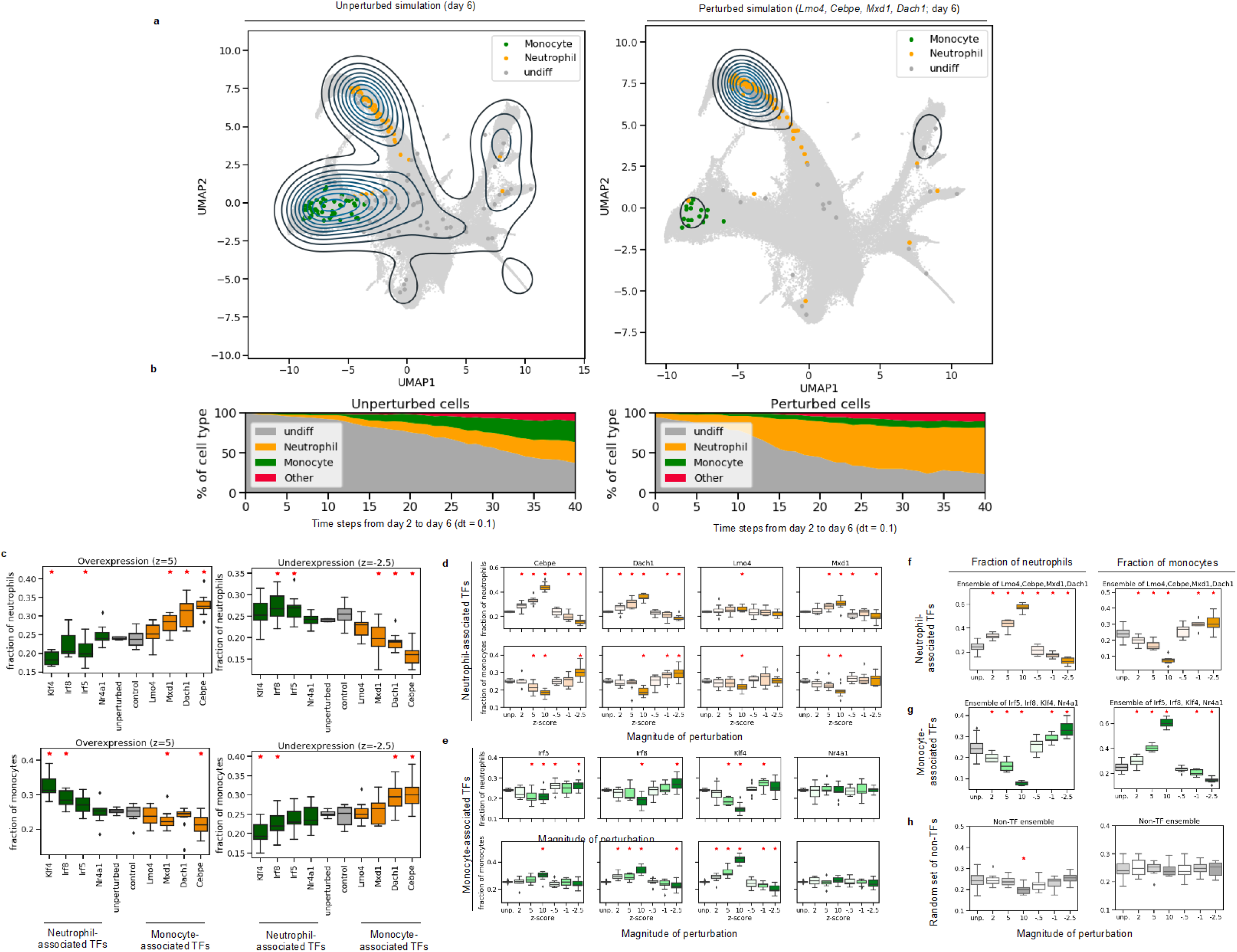
*in silico* perturbations of hematopoiesis results in expected shifts in fate distribution. (a) The distribution of cells at the final time-point generated by the model initialized with unperturbed cells (left) and cells with perturbations of *Lmo4, Cebpe, Mxd1, and Dach1* upregulated (z = 10) during neutrophil differentiation (right). (b) Proportions of generated cell types from day 2 to 6 initialized with unperturbed cells (left) and cells with perturbations of transcription factors upregulated during neutrophil differentiation (right). (c) Fraction of neutrophil and monocyte cells at final time point with single-gene perturbations. Transcription factors involved in monocyte development are indicated in green, while transcription factors involved in neutrophil development are indicated in orange. Control genes (in grey) indicate experiments when perturbing genes from a random set of non-TFs as in (h) (d-e) Individual genetic perturbations made to transcription factors involved in neutrophil and monocyte differentiation have an increased effect at higher dosages. (f-g) Ensemble perturbations of transcription factors involved in neutrophil (orange) and monocyte (green) differentiation have an ever stronger effect. (h) Ensemble random perturbations of non-transcription factors (grey) have no trend in cell type proportion change. (c-f) Red asterisks indicate independent t-test at p < 0.05 with respect to the unperturbed simulations over 10 randomly sampled starting populations.

**Figure 5.**
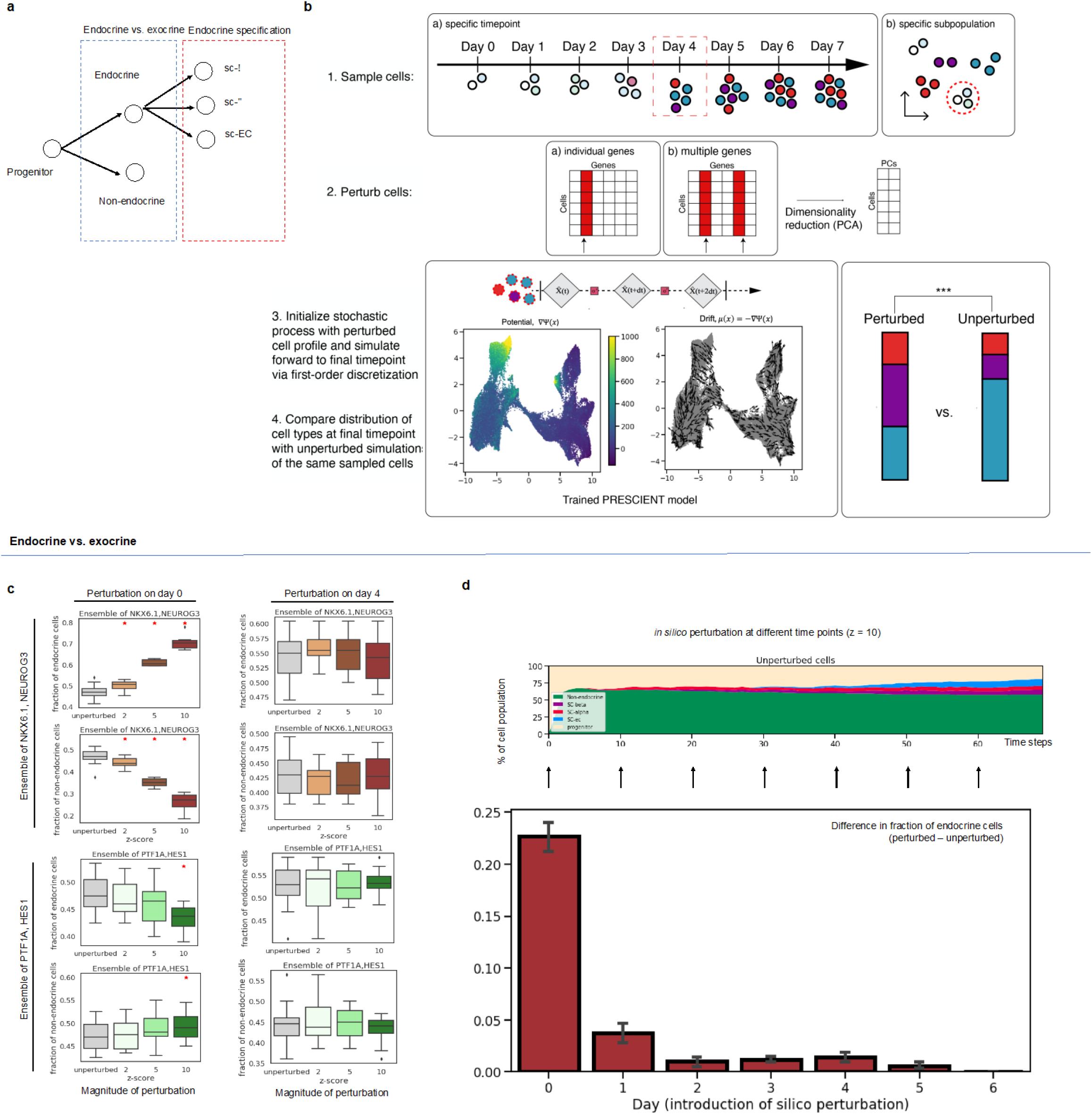
PRESCIENT predicts expected temporal dynamics of *in silico* perturbations of the endocrine/exocrine axis. (a) Simplified model of *in vitro* endocrine induction. Progenitor cells first bifurcate along the endocrine-exocrine axis, and then are specified into more specific pancreatic cell fates. (b) Schematic of steps for different perturbational experiments. First, cells are sampled from either a specific timepoint in the scRNA-seq timecourse or by a specific subtype label. Next, either individual genes or ensembles of genes are perturbed by setting target genes to a higher or lower z-score in scaled gene expression space. This profile is then reduced to principal components. These perturbed cells are then used to initialize a simulation of an already trained PRESCIENT stochastic process, which is stepped forward to the final timepoint. Finally, a pre-trained ANN classifier is used to evaluate the cell type distribution at the final time-step of the simulation. This distribution is then compared to an unperturbed simulation via a paired t-test. (c) Final fractions of endocrine and exocrine cells as a result of ensembled perturbations of endocrine- and exocrine-associated TFs (top, bottom) on day 0 and day 4 (left, right) with increasing magnitude. Red asterisks indicate significance at independent t-test p < 0.05. (d) *in silico* perturbations are introduced at different time points to the corresponding unperturbed population. The different outcomes of the cell type of interest are then calculated as the difference in fraction at the final time point starting from the perturbed and unperturbed populations. (c-d) Results are reported over 10 randomly sampled starting populations.

#### Perturbation of transcription factors involved in the regulation of hematopoiesis

We hypothesized that a PRESCIENT model trained on the Weinreb et. al. dataset (2 layers of 400 units, seed 1, epoch 2500) should be able to recapitulate the effects on cell fate when perturbing transcription factors known to be involved in regulation of neutrophil or monocyte differentiation. The model predicts the outcome of each perturbation on all cell types represented in the dataset (Figure S2). In particular, we focus on a set of transcription factors previously identified to be potentially antagonistically correlated with either monocyte and neutrophil fate in progenitor cells by MetaCell analyses of a haematopoietic stem cell dataset, many of which are supported by existing literature. The same authors observed in CRISPR-seq experiments that Irf8 deficiency correlated with increase of neutrophils and blocked monocyte differentiation (Giladi et al., 2018). Existing literature also supports several of the transcription factors identified. For example, Feinberg et al. previously showed that forced expression of Klf4 in hematopoietic stem cells induces exclusive monocyte differentiation, while Klf4 deficiency increases granulocyte (of which neutrophils are the most abundant type) differentiation (Feinberg et al., 2007). As another example, Ly6C-monocytes have previously been found to be absent in Nrf4a1-/- mice (Hanna et al., 2011). With regards to transcription factors associated with neutrophil fate, CEBPE deficiency in -/- mice has similarly been shown to block maturation of granulocyte lineages (Yamanaka et al., 1997). Mutations in CEBPE are also one of the known genetic causes of specific granule deficiency, a rare disorder characterized by abnormal neutrophils (Serwas et al., 2018).

We first perturbed *Lmo4, Cebpe, Mxd1*, and *Dach1*, transcription factors previously identified to be involved in granulopoiesis, the production of mature neutrophils. These transcription factors were also present in the highly variable gene set from the Weinreb et. al. hematopoiesis time course (Figure 4c). To simulate over-expression, we set the scaled normalized expression of the target gene(s) to 5, while to simulate knock-down/out, we set the scaled normalized expression of the target gene(s) to −2.5. As expected, we observed that down-regulation of these TFs led to a relative decrease in the fraction of neutrophils while up-regulation of these TFs led to a relative increase in the fraction of neutrophils with respect to the unperturbed population (Figure 4c). We next perturbed transcription factors involved in monocyte development, including *Irf8, Irf5, Klf4*, and *Nr4a1*, and observed similar results (Figure 4c).

We next tested if different magnitudes of the perturbation had different effects. For overexpression, we tested z-score = 2, 5, 10, while for knock-down/out, we tested z-score = −0.5, −1, −2.5. In general, we found that increasing the magnitude of the perturbation resulted in larger changes in the relative fraction of neutrophils (Figure 4d-e). Furthermore, while individual perturbations resulted in a mixture of significant and non-significant changes in the final neutrophil populations, ensemble perturbations consistently resulted in significant changes (p<0.5) in the fraction of the neutrophil population (Figure 4f-g). We also tested multiple sets of randomly selected non-TFs to ensure the changes in final cell fate were not simply a result of perturbations causing random model changes. These randomly selected non-TFs do not result in an observed shift in neutrophil and monocyte fates in the final time point (Figure 4h), suggesting that our model is robust to random effects.

#### Perturbation of gene sets involved in the regulation of β-cell development induced at multiple timepoints and developmental stages

We next applied PRESCIENT to another well-characterized differentiation system, the production of pancreatic islet cell types *in vitro*. Recently, Veres et al. collected a 7 time-point scRNA-seq time-course dataset of *in vitro* endocrine induction using their human pancreatic islet differentiation protocol (Figure 5a, S3a-b, e-f). This dataset did not include lineage tracing measurements. Applying PRESCIENT to this dataset, we first show that we are able to recapitulate the expected effects of perturbations to TFs that are well-documented regulators of pancreatic islet formation and specification. We next illustrate the ability to introduce time-resolved perturbations by sampling and perturbing cells at different time points in the scRNA-seq dataset. Finally, we show that PRESCIENT can isolate the effect of perturbations on individual transition cell types along the axis of endocrine induction from pancreatic progenitor to terminal fate.

We first hypothesized that PRESCIENT should be able to recapitulate the effects on cell fate when perturbing transcription factors known to be involved in the regulation of endocrine induction and specification in the starting population (Figure 5a-b). In endocrine/exocrine induction, previous work has shown that NEUROG3 and NKX6 activation is associated with the endocrine lineage, while PTF1A and HES1 is associated with the exocrine lineage (Dassaye et al., 2016; Gradwohl et al., 2000; Johansson et al., 2007; van der Meulen and Huising, 2015). For example, (Jensen et al., 2000) showed that HES1 activation results in an enrichment of the exocrine cell lineage, while (Johansson et al., 2007) showed that overexpression of NEUROG3 results in general expansion of endocrine cells. Furthermore, PTF1A and NKX6 have been shown to act antagonistically (Schaefer and Peifer, 2019), as do NEUROG3 and HES1 (Jensen et al., 2000), when modulating the endocrine/exocrine fates. When introducing ensembled in silico perturbations of NEUROG3 and NKX6.1 at day 0, we observe that this results in an increase in endocrine cell types that scales with the magnitude of perturbations (p<0.05) with a reciprocal decrease in exocrine cell types. We also show that PTF1A and HES1 overexpression result in the opposite effect (Figure 5c). We observed corresponding decreases in endocrine and exocrine cell proportions with knock-downs of NEUROG3/NKX6.1 and PTF1A/HES1, respectively (Figure S5a-b)

We next simulated the effects of TFs involved in the specification of cell types post-endocrine induction, where ARX and PAX4 have previously been shown to have an antagonistic effect on the specification of endocrine precursors to α and β-cell fate respectively (Dassaye et al., 2016). These TFs have been shown to have a reciprocal repressive effect via direct interaction with their respective promoters. ARX-deficient mice show severe hypoglycemia and an increase in β-cell production (Collombat et al., 2003, 2005). In CRISPR-Cas9-induced homozygous PAX4 knock-out rabbits, severe hyperglycemia was observed as a result of a major decrease in insulin-producing β-cells and a corresponding increase in α-cells (Xu et al., 2018). Introducing computational overexpression of ARX (z=5,10) to progenitor cells at day 0 resulted in significant increases in the final proportion of α cells at the expense of β cells (Figure 6a). The same overexpression of PAX4 resulted in a significant increase in the final proportion of β cells (p<0.05) at the expense of α cells (Figure 6a). We observed similar results when performing underexpression experiments (Figure S4).

**Figure 6.**
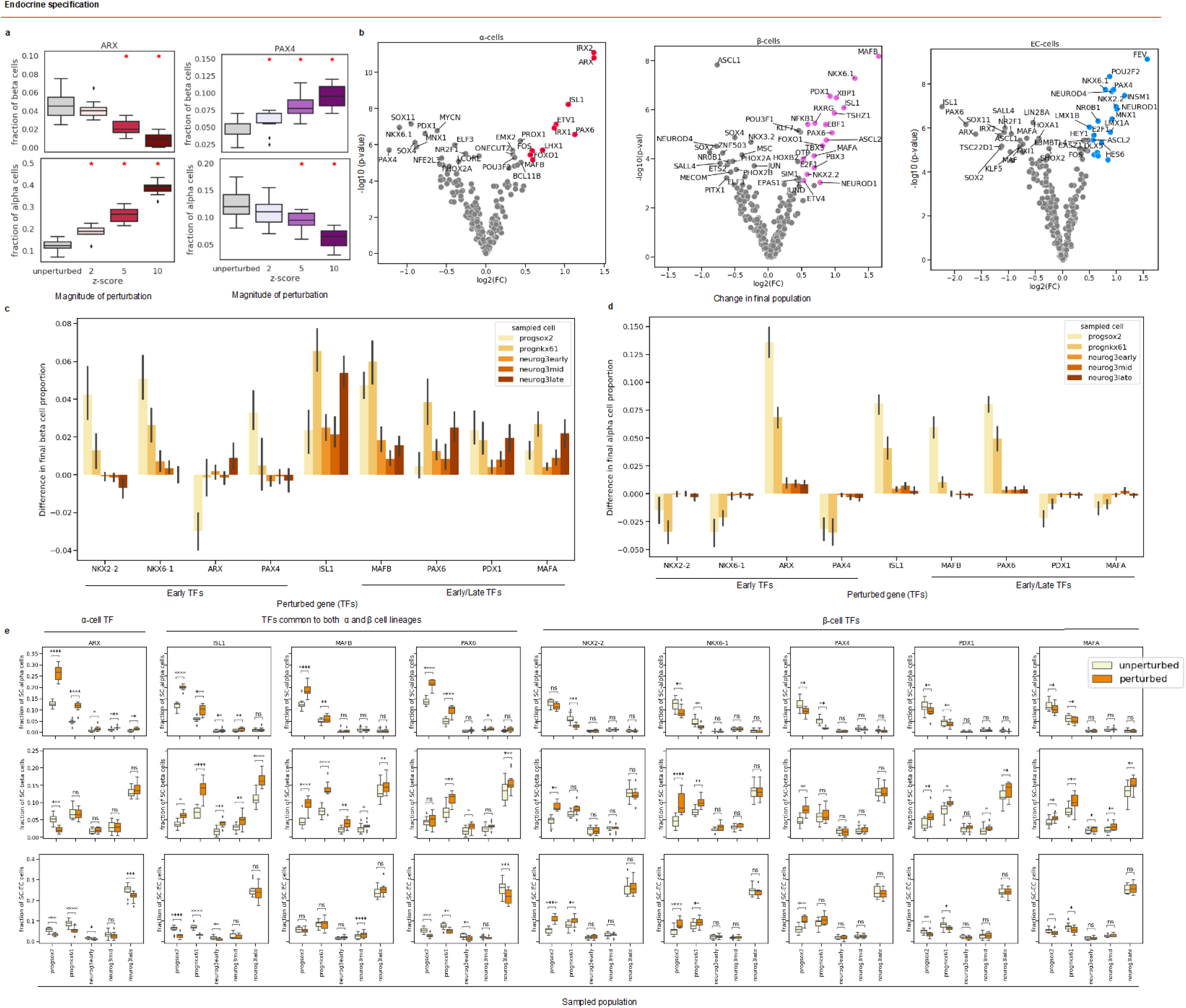
PRESCIENT predicts the outcome of transcription factor perturbations in a large perturbational screen as well as the different effects of perturbations in early progenitors vs. cells further along endocrine induction. (a) Final fractions of α and β cells when introducing perturbations to ARX and PAX4 in the starting cell population at day 0. Red asterisks indicate significance at independent t-test p < 0.05. (b) Perturbational screen of all the TFs in the highly variable gene set of *in vitro* β-cell differentiation. The x-axis is the log2 fold change (FC) of the final cell type fraction of target cell fraction between perturbed and unperturbed simulations. The y-axis is the −log10 p of a paired t-test between unperturbed and perturbed cell fractions at the final time step. Points are colored if they are a hit (p < 0.01 & log2(FC) > 0.5) and β-cell fractions are shown in purple, α-cells in red, and EC-cells in blue (c) Difference in final β-cell perturbations when introducing perturbations in different cell populations for different TFs (d) Difference in final α-cell perturbations when introducing perturbations in different cell populations for different TFs (e) Final fractions of α, β and EC cells when starting from different perturbed vs. unperturbed populations. Asterisks indicate significance at paired t-test p * < 0.05 ** < 0.01, *** < 0.001. (b-c) Results are reported over 10 randomly sampled starting populations.

To demonstrate the scale at which *in silico* experiments can be conducted, we next did a larger screen of 200+ TFs (Figure 6b). At p < 0.01 & log2(FC)>0.5, we found that this unbiased *in silico* screen identified 10 TFs (α specification increase) and 18 TFs (β specification increase) that altered cell fate. The computational screen identified several known fate-specific factors, such as ARX and IRX2 for α-cells (Baron et al., 2016; Collombat et al., 2003) and NKX2.2, NKX6.1, PAX4, PDX1, and MAFA for β-cells. NKX2.2 has previously been associated with repression of the α-cell fate, along with NKX6.1 and PDX1 (Gannon et al., 2008; Papizan et al., 2011; Schaffer et al., 2010, 2013). MAFA expression is generally siloed to β-cells, and in MAFA-deficient mice about two-thirds of the β-cell population is lost (Artner et al., 2010). This screen also identified factors common to both alpha and beta-cell lineages, including MAFB, PAX6, and ISL1. PAX6 has previously been reported to be associated with both α and β-cell fate, but Pax6 mutant mice show a more significant decrease in α-cells, which we also observe. In addition, we identified 19 TFs for EC specification (pval < 0.01 & log2(FC)>0.5), showing how PRESCIENT can be used to generate hypotheses for CRISPR genetic screens (Figure 6b).

We next asked if PRESCIENT could recapitulate the known timing of TFs during endocrine induction. To this end, we performed experiments in which we introduced perturbations to the same endocrine/exocrine axis TFs described before at multiple timepoints (days 0-6) and then simulated forwards in time to the final time point with a *dt* (step size parameter) of 0.1. Endocrine/exocrine induction is known to occur early in the time course, and the model results corroborate this as early perturbations to NKX6.1 and NEUROG3 (z=10) result in an increase in endocrine cells, but this effect diminishes with perturbations induced at later time points (Figure. 5d).

PRESCIENT also enables perturbations of cells sampled from different starting cell populations along a given differentiation trajectory. To demonstrate this, we introduced perturbations of selected TFs from the larger screen to cells sampled from different stages (early to late) of the endocrine induction pathway as labeled in Veres et al.: SOX2+ progenitors (*progsox2*), NKX61+ progenitors (*prognkx61*), and cells with early/middle/late NEUROG3 signatures (Figure 6c-e); *neurog3early, neurog3mid, neurog3late*, respectively). NKX2.2, NKX6.1, PAX4, and ARX are found early in endocrine specification and are often the first signal of the production of specific terminal endocrine cell fates (van der Meulen and Huising, 2015). We found that perturbations of these TFs in pancreatic progenitors result in a significant increase in final β-cell proportion (p<0.05) and this effect is minimal in cells further along the endocrine induction pathway (Figure 6c, e). In contrast, PDX1 has been shown to continue to promote β-cell neogenesis late into the endocrine induction pathway. Forced expression of PDX1 in NEUROG3^+^ cells increased the ratio of β-cells to α-cells in mouse embryos and adult mice (Yang et al., 2011). We show that computational perturbations to PDX1 increase the fraction of β-cells when introduced in progenitor cells and neurog3late cells (Figure 6c, e), recapitulating the multiple contexts in which perturbations to PDX1 can modulate endocrine fate. Similarly, MAFA and MAFB are known to continue to modulate fate late into endocrine induction (Artner et al., 2010; Hang and Stein, 2011), and we show that perturbations to both MAFA and MAFB result in increases in β-cell proportion when introduced to both early pancreatic progenitors and cells in late endocrine induction (Figure 6c,e). While ISL1 has been previously reported to stimulate both α and β-cell generation, the timing of ISL1 activation is less well characterized. Our model suggests that ISL1 has a role in endocrine specification in both early pancreatic progenitor phase and late in the endocrine induction pathway (Figure 6c,e). Finally, we observed that modulation of α-cell fate largely occurs via perturbations induced in the pancreatic progenitor phase of the stage 5 differentiation protocol, with diminished effects of all endocrine specification TFs in *neurog3early, neurog3mid*, and *neurog3late* (Figure 6d,e). This could suggest that α-cell determination occurs early in the stage 5 differentiation protocol. We observe similar outcomes with *in silico knockdown* experiments (Figure S5) as well as no effect of perturbations to randomly sampled non-TF genes that are not involved in apoptotic/proliferative signatures (Figure S4c).

### Discussion

PRESCIENT is a generative modeling framework for learning potential landscapes from population-level time series scRNA-seq data. We build upon previous work in two important ways. First, we used deep learning to approximate complex potential functions that underlie development. Second, we incorporated cell proliferation into the modeling framework via weighting of cells in the loss function. We validated PRESCIENT on a recent experimental lineage tracing dataset that provides gold standard trajectories for the mouse hematopoietic lineage, and show that it outperforms other existing fate prediction methods on the task of predicting the final cell fate distribution of a given progenitor cell.

We found that the incorporation of cell proliferation rates greatly improved the performance of PRESCIENT cell fate prediction on a gold-standard lineage tracing dataset. Cell proliferation is well-known to be an important part of cellular differentiation but is often neglected when modeling differentiation. When lineage tracing data is unavailable we estimate cell proliferation rates in an approach similar to that previously described by Schiebinger et al. Incorporating iterative updating of cell proliferation rates into our framework in the future would reduce reliance on our gene-expression-based estimates (Schiebinger et al., 2019).

PRESCIENT marks a departure from the predominant methods for analyzing scRNA-seq studies of cellular differentiation. Computational methods for lineage inference have been dominated by pseudo-time approaches that do not attempt to model the stochastic or dynamic nature of cell fate determination. More recent fate prediction methods such as Waddington-OT and Population Balance Analysis either summarize observations of the emergent process or suffer from modeling limitations (Schiebinger et al., 2019; Weinreb et al., 2018). However, we have demonstrated an important predictive advantage to fully generational models that seek to describe the underlying differentiation landscape. After the model has learned the landscape, it can generate trajectories for unseen data points.

Generative models can hence be interrogated to propose hypotheses for possible perturbations. We show that inducing perturbations in well-studied regulatory genes of hematopoiesis and β-cell differentiation result in expected changes in fate outcome. We also show that we can introduce complex perturbations consisting of multiple genes, or at different time points or in different starting populations. The ability to study how these different factors interact enables large computational experiments that can help limit the number of *in vitro* experiments needed to achieve a desired cell fate, especially when considering combinatorial perturbations and timing of perturbations in a differentiation process. Using PRESCIENT to identify genetic targets of CRISPR-screens and small molecules can help aid the design of new directed differentiation and reprogramming protocols as well as fine-tuning existing approaches to improve cell fraction outcomes. We show an example of this type of unbiased, large-scale computational perturbational screen for *in vitro* β-cell differentiation in which we perturbed 200+ TFs and identified target genes that could cause significant shifts in β-cell, α-cell, and EC-cell fates. While this work was limited to TFs to show the utility of the method, non-TF targets, such as signaling pathway effectors, can also be tested using PRESCIENT. However, it is important to note that the model is subject to constraints and assumptions. For example, learning a model requires that the final time point of the dataset is at steady-state, and model performance improves with more densely sampled time-points. There also remain challenges to confidently suggest gene sets for experimental perturbation. One problem is that information is lost about individual genes when transforming data into PCA space, and lowly-expressed genes important to cell fate decisions may be dropped altogether in the scRNA-seq data. Another is that the association of certain genes with specific cell fates does not necessarily imply causality.

Future extensions of generative modeling can accommodate other forms of data. For example, with lineage tracing data a model’s objective function can be modified to maximize the likelihood of observing individual trajectories. Data on RNA velocity (La Manno et al., 2018) can be used to constrain the drift function directly. We expect that constraining the model with additional sources of information should improve the quality of the underlying landscape inferred.

## Acknowledgements

We thank Jennifer Hammelman and Ernest Fraenkel for helpful discussion and comments. We acknowledge funding from NIH grants 5 R01 NS109217 and 5 R01 HG008363 (D.K.G) and the National Science Scholarship (PhD) from the Agency for Science, Technology and Research Graduate Academy (G.Y.).

## Author contributions

Conceptualization, Methodology, Software, Validation, Formal Analysis, Investigation, Data Curation, Writing, Visualization: G.Y., S.S.; Conceptualization, Writing, Formal Analysis, Supervision, Funding Acquisition: D.K.G.

## Methods

### Method details

#### Identifying the latent dynamics of cellular differentiation

Following Hashimoto et al. (Hashimoto et al., 2016), we model cellular differentiation as a diffusion process *X*(*t*) given by the stochastic differential equation

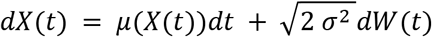

where *X*(*t*)represents the *k*-dimensional state of a cell at time *t*, *μ*(*X*(*t*)) is a drift term representing the force acting on a cell given its state, and *W*(*t*)corresponds to unit Brownian motion. In particular, the drift function is defined to be the negative of the gradient of a potential function, *μ*(*x*) = −Δ*Ψ*(*x*), such that intuitively, the potential function *Ψ*(*x*) can be thought of as inducing a gradient field driving cells from regions of high potential to low potential. Within the conceptual framework of Waddington’s epigenetic landscape, this potential function corresponds to the height of the landscape. This process can be simulated via first-order time discretization

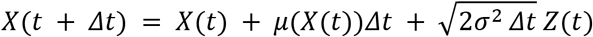

where *Z*(*t*) are i.i.d. standard Gaussians. This converges to the diffusion process as Δ*t* → 0.

We define the marginal distribution at time *t* to be *ρ*(*x*, *t*) = *P*(*X*(*t*) = *x*). The inference task identifies the potential function *Ψ*(*x*), and hence the underlying drift function *μ*(*x*), given only samples from the marginal distribution {*x*(*t*)_*i*_ ~ *ρ*(*x*, *t*) | *i* ∈ {1… *m_t_*}, *t* ∈ {1.. *n*}, where *m_t_* is the number of cells sampled at time *t* and *n* is the number of time points where data was observed. In practice, this data corresponds to gene expression profiles of cells sampled over the course of a time-series experiment.

Inference proceeds by finding the potential function *Ψ* in a family of functions K that minimizes the objective

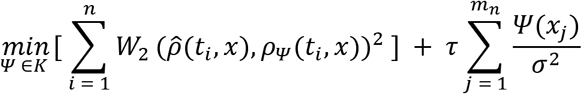

where 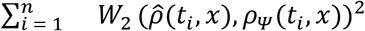 is the Wasserstein distance between the empirically observed distribution *ρ*(*t_i_*, *x*) and the predicted distribution for a candidate potential function *ρ_Ψ_*(*t_i_*, *x*), and *τ* is a parameter controlling the strength of the entropic regularizer. We refer the reader to Hashimoto et al. for derivation of this objective function.

### Incorporating cell proliferation

Using notation from Feydy et al. (Feydy et al., 2019), computing the Wasserstein distance involves solving the optimization problem

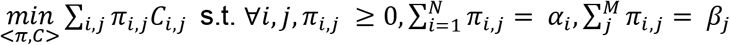

where *π_i,j_* is an optimal transport plan mapping points from the source measure to the target measure, *C_i,j_* = ||*x_i_* – *y_j_*||^2^ is the squared euclidean distance between point *x_i_* and point *y_j_*, and *α_i_* and *β_j_* are positive weights associated with the samples *x_i_* and *y_j_*. Previous models minimizing the Wasserstein loss as described by Hashimoto et al. as well as Schiebinger et al. set the weights *α_i_* and *β_j_* to be constant, hence assuming uniform sampling over the source and target densities. We propose to incorporate cell proliferation into the modeling framework simply by settings *α_i_* to the number of descendants the cell *x_i_* is expected to have, where in this case *x_i_* corresponds to samples from the predicted population. Weights associated with the empirical population *β_j_* remain constant. Note that in contrast, Schiebinger et al. incorporate cell proliferation by instead using the cell proliferation rates to weaken the marginal constraints, hence reframing the problem as unbalanced optimal transport.

Where lineage tracing data is available, we define the number of descendants a cell is expected to have at time *t* to be the number of cells sharing a particular clonal lineage barcode at *t* divided by the number of cells sharing that clonal lineage barcode at the current time. A pseudocount of 1 is added to allow for dropout.

In the absence of lineage tracing data, we estimate the number of descendants using a modified approach from Schiebinger et al.. Let *n* be the number of descendants, *b* be the birth rate, *d* be the death rate, and *g* be growth. Then using a birth-death process,

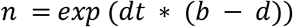

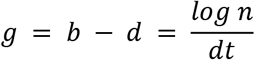

To estimate the birth and death scores (*s_b_*, *s_d_*), we first calculate the mean of the z-scores of genes annotated to birth (KEGG_CELL_CYCLE) and death (KEGG_APOPTOSIS). We use these alternative annotations as many of the genes in the original gene sets proposed by Schiebinger et al. were not present in the set of variable genes in the Weinreb et al. datasets. Then, following Weinreb et al., we smooth these scores over the cells using an iterative procedure:

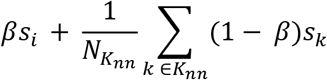

Where *s_i_* is the score for a given cell *i* at the current iteration, *s_k_* are the scores for the *k* = 20 neighbors of the cell *i* computed on euclidean distance in PCA space, and *β* = 0.1. This smoothing procedure is performed over 5 iterations.

Then, the birth and death scores are fit to logistic functions to obtain the birth and death rates,

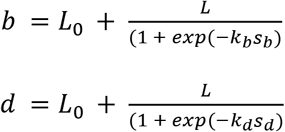

To simplify the fitting procedure, we reasoned that we might expect −*s_min_* to be close to 1 to flatten outliers. Hence, we set *k* to be 0.001 = *exp* (−*k* – *s_min_*). Then we set *L*_0_ and *L* based on the expected minimum and maximum growth rates. For the Weinreb et al. dataset, we set *L*_0_ = 0.3 and *L* = 1.2, and found the minimum and maximum number of descendants to be similar to that observed in the clonal lineage tracing data at days 4 and 6. We used the same procedure for estimating growth rates on the Veres et al. dataset.

#### Model implementation and optimization

We use deep learning methods to learn the parameters of the potential function *Ψ*(*x*). Automatic differentiation can be used to evaluate the drift function *μ*(*x*) = −*ΔΨ*(*x*) without having to derive the analytical form. Then, by retaining the computation graph, losses can be calculated with respect to the drift function while gradients are accumulated with respect to the potential function. For all models, we set dt and sd to be 0.1.

Optimization of *Ψ*(*x*) was performed using the Adam optimizer with a batch size of 1/10th the size of the training set. Models were pre-trained as previously described by Hashimoto et al. by first optimizing only the entropic regularizer via contrastive divergence using stochastic gradient descent with a learning rate of 1e-9. Then, training proceeds at the learning rate described below, with a scheduler that multiplied the learning rate by 0.9 every 100 iterations, and using gradient clipping with a max norm of 0.1. The Wasserstein error is approximated using a multi-scale Sinkhorn algorithm as implemented by the GeomLoss library (v0.2.3), with a scaling of 0.7 and a blur of 0.1. The GeomLoss library allows for efficient, stable computation of gradients that bypasses naive backpropagation (Feydy et al., 2019).

All models were implemented in PyTorch 1.4.0 (Paszke et al., 2019). Models were trained on either Titan RTX or GeForce RTX 2080 Ti GPUs.

All nearest-neighbors calculations (for eg. for cell-type classification) were calculated using the approximate nearest-neighbors library annoy (https://github.com/spotify/annoy).

### Quantification and statistical analysis

#### Preprocessing of existing scRNA-seq datasets

Data for the Weinreb et al. experiments was downloaded from https://github.com/AllonKleinLab/paper-data/blob/master/Lineage_tracing_on_transcriptional_landscapes_links_state_to_fate_during_differentiation/README.md (Weinreb et al., 2020). Normalized gene expression for genes annotated as variable was scaled and projected to 50 dimensions via PCA, which was then used as input to the modeling framework. For experiments evaluating the model on the held-out time point, preprocessing was fit to only the training set consisting of days 2 and 6, and then used to transform all data including day 4. For all other experiments, preprocessing was fit to all data across time points. All visualizations using umap were fit with 30 neighbors.

Data for the Veres et al. experiments was downloaded from GEO (GSE114412) (Veres et al., 2019). Raw counts were first pre-processed using the standard Seurat pipeline (Butler et al., 2018) to obtain normalized counts. For feature selection, genes were first filtered for those observed in at least 10 cells. Then, the ‘FindVariableFeatures’ function was used to identify the top 2500 most variable genes. Scaled gene expression was then computed as for the Weinreb et al. dataset. For projection into PCA space (30 PCs), the variable gene set was filtered again to remove genes correlated with TOP2A (r > 0.15), as described by Veres et al. This was used as input to the modeling framework, and for visualization via UMAP. For estimation of proliferation rates, the full variable gene set was used (Figure S3a-b, e-f).

#### Comparing model architectures on recovering a held-out time point

Architectures were chosen such that the 2 layer models had a corresponding 1 layer model with a roughly equivalent total number of parameters. The smaller models (1 layer of 1000 units, 2 layers of 200 units) had ~50k parameters and were trained using Adam with a learning rate of 0.01. The larger models (1 layer of 4000 units, 2 layers of 400 units) had ~200k parameters and were trained with a learning rate of 0.005, as these models sometimes diverged when trained with a learning rate of 0.01 (Figure S1a). All models were trained for 2500 epochs. Softplus was used as the activation function for all models.

To evaluate the models at a given time point, 10000 cells were sampled at day 2 with replacement according to the expected cell proliferation rate. Then, the model was used to sample a single trajectory for each of the sampled cells until the time point under evaluation. The Wasserstein distance was then computed between the simulated cell population and the empirically observed cell population. Models were evaluated at day 4 (training) and day 6 (held-out, testing) every 100 training epochs. The testing distance is reported for the epoch with the lowest training error.

As a baseline, we compared to Waddington-OT (WOT), which uses a similar optimal transport formulation but lacks an explicit parametric form (Schiebinger et al., 2019). WOT enables recovery of a held-out time point via linear interpolation using transport maps built between sets of cells in early and late time points. To run WOT, we used python code available on GitHub (https://github.com/broadinstitute/wot). The input to WOT is a set of time-point labeled gene expression profiles and growth rates and the output is an optimal transport map. The optimal transport map was built with the full set of cells with lineage barcodes from day 2 (n=4,638 cells) to day 6 (n=29,679 cells). The ground-truth proliferation rates derived from clonal expansion of this set of cells from day 2 to 6 were provided to WOT and three growth iterations were permitted. The parameters for building the optimal transport map were as follows: λ_1_=1, λ_2_=50, ε=1, τ=10000. With the transport map built between day 2 and day 6, 10,000 cells at day 4 were interpolated using the interpolate_with_ot() function from WOT. This maps a point at the midpoint of each of the pairs in the optimal transport map. The testing distance of these interpolated points from the observed day 4 cells was computed as reported above.

#### Predicting clonal fate bias

To predict clonal fate bias, models with 2 layers of 400 units and a regularization strength of tau = 1e-6 were first trained on data from all three time points. Models were fit on three sets of data; (a) the subset of cells for which lineage tracing data is available, (b) the subset of cells for which no lineage tracing data is available, and (c) all cells. Models were trained for 2500 epochs and evaluated every 500 epochs. Then, cells were simulated until the final time point via the first-order discretization as parameterized by the trained model.

We evaluated the clonal fate bias metric described by Weinreb et al. on the test set also defined in their paper. The approximate nearest neighbor (ANN) classifier that we used to classify cells as Neutrophil, Monocyte or other at the final time point was fit with 10 trees, 20 neighbors and using Euclidean distance in PCA space (50 pcs). The model was first fit on a random 80% split of the data. When evaluated on the held-out 20% test split, the model achieved a macro-average f1-score of 0.98. Splits were stratified by cell type. The model was then re-fit to the full dataset. After classification, the clonal fate bias was then computed as the number of neutrophils divided by the total number of monocytes and neutrophils. Since the model did not always predict any cell within the 2000 sampled trajectories to be a monocyte or neutrophil, we also added a pseudocount of 1. In those cases, clonal fate bias would hence be 0.5.

We observed variation in predictions made by models at each epoch (Figure S1e). Since no validation set is available to formulate a stopping criteria, we chose to ensemble predictions over the last 5 epochs evaluated (i.e. epoch 2100, 2200, 2300, 2400, 2500) by taking the mean across the estimated clonal fate bias across those epochs.

In most cases, performance metrics for WOT, PBA and FateID were calculated using the predictions already pre-computed and made available by Weinreb et al. (Weinreb et al., 2020)

#### Introducing and evaluating *in silico* perturbations

Perturbation experiments were performed similarly to the clonal fate bias experiments, except using perturbed cells as input to the first-order discretization. Generally, perturbations were introduced *in silico* by setting the scaled normalized expression of target genes to z-score values less than 0 for knockdowns and greater than 0 for overexpression. The resulting perturbed gene expression profile was then transformed via PCA into lower-dimensional space for input to forward simulation of the trained model. In this section, we describe perturbational experiments performed by leveraging PRESCIENT models trained on longitudinal scRNA-seq datasets characterizing *in vitro* hematopoiesis and *in vitro* beta-cell differentiation.

For the *in vitro* hematopoiesis dataset (Weinreb et al., 2020), the trained PRESCIENT model used for predicting the effect of *in silico* perturbations to this dataset was seed 1 and epoch 2500 of a model trained with a neural network architecture of 2 layers of 400 units. For all experiments, 200 undifferentiated cells (annotations from Weinreb *et al.*) were randomly sampled from day 2 weighted by KEGG-derived growth rate estimates, resulting in biased sampling for actively proliferative cells. These sampled cells were simulated forward 40 steps with a dt of 0.1 to the final time point (day 6). This process was repeated 10 times with random initializations of both unperturbed cells and perturbed cells. Cells at the final time point were then classified using the same ANN classifier used for the clonal fate bias experiments. Relevant TFs for the target cell fate were identified by searching the literature for studies that had experimentally verified sets of TFs involved in early cell fate decisions by progenitor populations and the highly variable feature set was filtered for these TFs. Perturbations were focused on monocytes and neutrophils due to the availability of experimentally correlated or confirmed perturbations for these two cell types and the focus of neutrophil/monocyte fate in the Weinreb *et al*. lineage tracing data. Neutrophil associated transcription factors were *Lmo4, Cebpe, Mxd1*, and *Dach1*. Monocyte associated transcription factors were *Irf8, Irf5, Klf4*, and *Nr4a1*. First, the effect of perturbations to individual genes were tested by perturbing each target gene with a z-score of −2.5 for knockdown and 5 for overexpression. To test if there was a significant shift in neutrophil/monocyte cell fractions at the final time point, Welch’s independent t-tests were performed between unperturbed simulations and perturbed simulations for each target TF. Next, the effect of perturbational magnitude was evaluated by introducing perturbations of −2.5, −1, −0.5, 2, 5, and 10 to each target gene individually. The same statistical test was performed in comparison to unperturbed simulations. Next, we tested the effect of ensembled perturbations to cell fate outcome by perturbing sets of TFs with z-scores of −2.5, −1, −0.5, 2, 5, and 10. As a control, the ensembled perturbation was repeated with following randomly selected non-TF control genes: *Gch1, Pfn2, Dhrs2, Traf1, Lrrk2, Lgmn, Il13*, and *Sgk1*. The same statistical test was performed in comparison to unperturbed simulations.

For the *in vitro* beta-cell differentiation dataset (Veres et al., 2019), we trained PRESCIENT models with 2 layers of either 200 or 400 units with a tau of 1e-6. A learning rate of 0.001 was used to prevent model divergence. All other parameters were as in the models trained for clonal fate bias prediction in the Weinreb et al. dataset. The trained PRESCIENT model used for predicting the effect of *in silico* perturbations to this dataset was seed 1 and epoch 1500 of a model trained with 2 layers of 400 units. The 400 unit model was chosen as it achieved a lower training distance, and epoch 1500 was chosen based on the training curves to prevent overfitting, since we observed that training performance had appeared to have plateaued by that epoch (Figure S3c-d). As with the Weinreb et al. dataset, an ANN classifier with 10 trees, 20 neighbors, and using Euclidean distance in PCA space (30 PCs) was trained to predict cell type on a training set of 80%. On a randomly held-out test set of 20%, the model achieved a macro-average f1 score of 0.939 when discriminating between sc-α, sc-β, sc-EC and other cells. The ANN classifier was then refit on the full dataset. For time point sampling experiments, 200 cells were sampled from each time point weighted by KEGG-derived growth rates, based on metadata from Veres *et al*. Cells were iteratively sampled from days 0-6 and simulated forward with a *dt* (step size parameter of 0.1) to the final time point (day 7) under both unperturbed and perturbed conditions. Perturbations (z=-2.5, −1, −0.5, 2, 5, 10) were introduced to sampled cells from each timepoint. To test if there was a significant shift in neutrophil/monocyte cell fractions at the final time point, Welch’s independent t-tests were performed between unperturbed simulations and perturbed simulations for each target TF. For cell type subpopulation experiments, 200 cells were randomly sampled from the SOX2+ progenitor, NKX6.1+ progenitor, and NEUROG3 early/mid/late, weighted by KEGG-derived growth rates. For the screen of 200+ TFs, cells were first simulated without perturbations and the same cells were simulated with perturbations (z=10) of each TF. This was repeated with 10 random initializations. For each TF, to test for a significant change in final cell type fractions, paired t-tests were conducted between the unperturbed and perturbed simulations.

### Resource availability

#### Data and code availability

All code as well as preprocessed data and trained models are available at https://github.com/gifford-lab/prescient

