## Supplemental Information for "Generative modeling of single-cell population time series for inferring cell differentiation landscapes"

Grace Hui Ting Yeo, Sachit D. Saksena, David K. Gifford

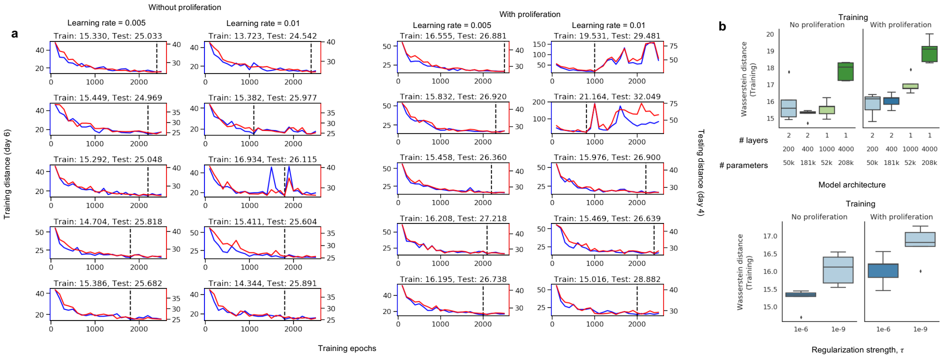


c

| Model | No proliferation 1e-3 | No proliferation 1e-6 | No proliferation 1e-9 | With proliferation 1e-3 | With proliferation 1e-6 | With proliferation 1e-9 |
|---|---|---|---|---|---|---|
| | | | Training (Day 6) | | | |
| 1 × 1000 | 60.53 ± 0.94 | 15.57 ± 0.48 | 16.61 ± 0.76 | 63.61 ± 0.43 | 16.99 ± 0.54 | 17.51 ± 0.74 |
| 1 × 4000 | 68.62 ± 1.49 | 17.82 ± 0.55 | 18.82 ± 0.59 | 71.45 ± 0.61 | 19.00 ± 0.71 | 19.90 ± 0.51 |
| 2 × 200 | 31.66 ± 9.18 | **15.88 ± 1.14** | 16.33 ± 0.63 | 29.91 ± 6.71 | **15.82 ± 0.67** | 17.17 ± 0.52 |
| 2 × 400 | 25.33 ± 3.21 | **15.23 ± 0.30** | 16.06 ± 0.44 | 27.45 ± 6.41 | **16.05 ± 0.42** | 16.78 ± 0.49 |
| | | | Testing (Day 4) | | | |
| 1 × 1000 | 49.76 ± 0.23 | 26.91 ± 0.48 | 28.50 ± 0.73 | 53.56 ± 0.90 | 28.11 ± 0.42 | 29.41 ± 0.44 |
| 1 × 4000 | 50.60 ± 0.60 | 28.09 ± 0.33 | 29.46 ± 0.24 | 54.19 ± 0.41 | 29.73 ± 0.52 | 31.27 ± 0.32 |
| 2 × 200 | 80.13 ± 8.99 | **25.78 ± 0.91** | 27.40 ± 1.03 | 89.11 ± 11.20 | **27.11 ± 0.09** | 28.23 ± 0.94 |
| 2 × 400 | 89.71 ± 9.35 | **25.31 ± 0.41** | 26.30 ± 0.83 | 117.31 ± 27.92 | **26.82 ± 0.31** | 27.64 ± 0.51 |

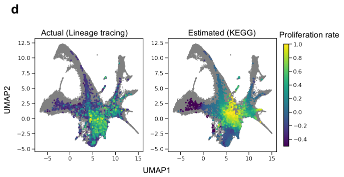


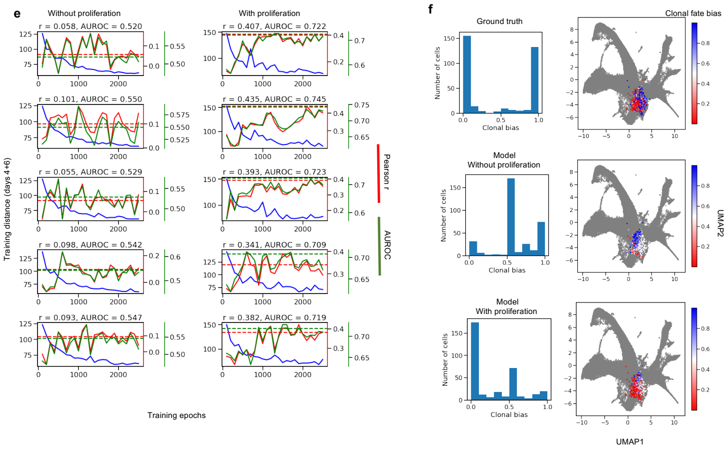


Figure S1: **Training and testing performance on time point recovery and clonal fate bias prediction tasks on Weinreb et al. lineage tracing dataset.**
(a-c) Results for time point recovery task (a) Training (blue, left axis) and testing (red, right axis) performance across training epochs for 2 layer 400 unit models with and without proliferation and with different learning rates across 5 seeds. Training and testing performance is reported for best training epoch, as indicated by the vertical dashed line (b-c) Testing performance for models of different complexity (above, table) and with different regularization strengths (below, table) for 5 seeds (d)Visualization of actual and estimated proliferation on UMAP (e-f) Results for clonal fate bias prediction task (e) Training (blue, left axis) and testing Pearson r (red, first right axis) and AUROC (green, second right axis) across training epochs for 2-layer 400 unit models with and without proliferation. Ensembled testing performance is reported and indicated as horizontal dashed lines (f) Distribution of ground truth and predicted clonal fate bias across testing cells as a histogram (left) and visualized on the UMAP (right).

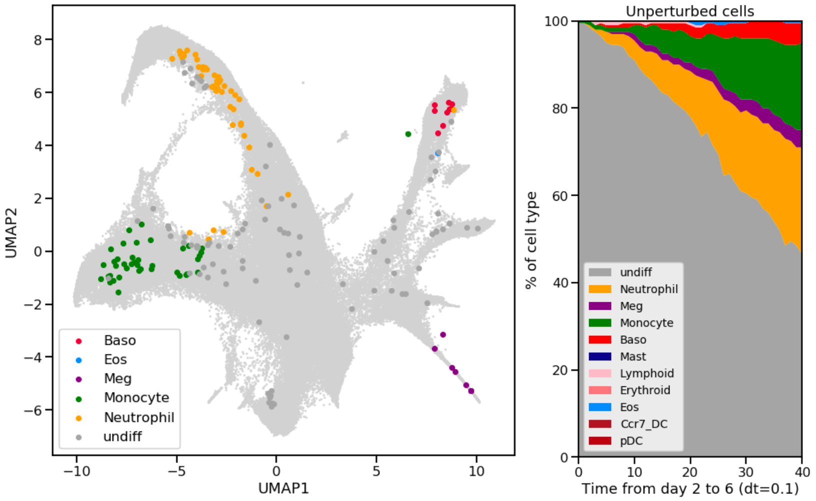


Figure S2: **Distribution of cell types at the final time point (left) and across training steps until final time point (right) for unperturbed simulations of in vitro hematopoietic differentiation.**

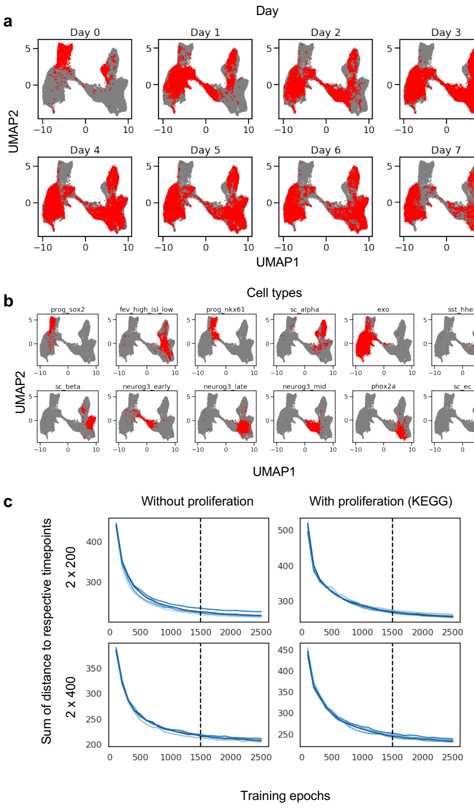


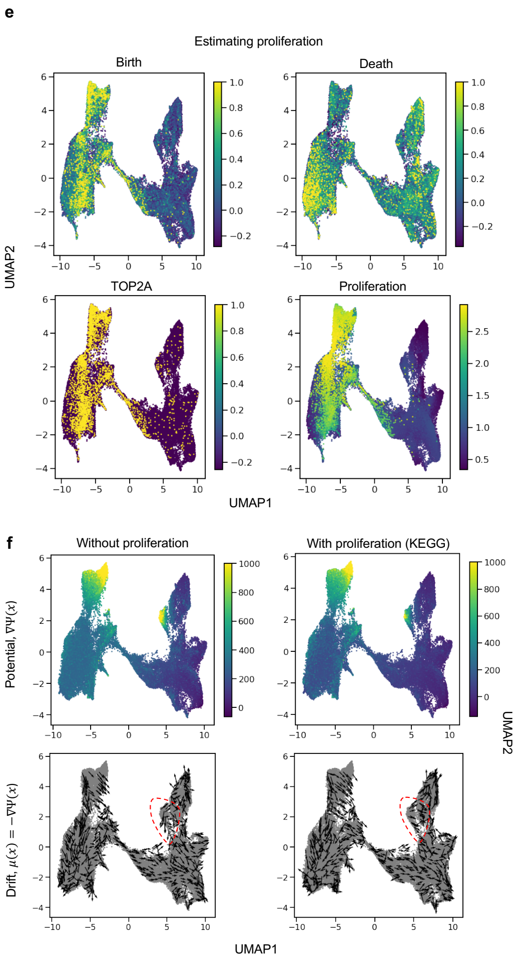


**d**

| Time point | No proliferation 2 × 200 | No proliferation 2 × 400 | With proliferation (KEGG) 2 × 200 | With proliferation (KEGG) 2 × 400 |
|---|---|---|---|---|
| 1 | 37.27 ± 0.78 | 35.19 ± 0.53 | 47.70 ± 1.28 | 44.51 ± 0.98 |
| 2 | 36.12 ± 0.68 | 34.45 ± 0.69 | 45.53 ± 1.28 | 41.04 ± 1.20 |
| 3 | 27.82 ± 0.65 | 26.87 ± 0.57 | 33.01 ± 0.65 | 30.66 ± 0.45 |
| 4 | 30.46 ± 1.12 | 28.63 ± 0.73 | 37.39 ± 0.69 | 33.46 ± 0.65 |
| 5 | 30.20 ± 1.08 | 28.46 ± 0.44 | 35.57 ± 0.76 | 31.90 ± 0.56 |
| 6 | 30.83 ± 1.10 | 30.17 ± 0.23 | 34.06 ± 0.48 | 31.40 ± 0.44 |
| 7 | 32.93 ± 0.77 | 32.27 ± 0.37 | 36.60 ± 0.45 | 33.81 ± 0.61 |

Figure S3: **Preprocessing and model fitting on Veres et al. dataset.** (a-b) Visualization of cells at different time points (a) or annotated as different cell types (b) (c-d) Training performance across 5 seed with and without proliferation for 2 2-layer models across training epochs and summed over time points (c) and for individual time points at epoch 1500 (d). The vertical dashed line in (c) also indicates epoch 1500. (e) Visualization of birth, death and proliferation rates estimated via KEGG, as well as scaled expression of TOP2A (f) Visualization of drift and potential learned by example 2 layer, 400 unit evaluated at epoch 1500. The red circles indicate qualitative differences in drift and potential between models fit with and without proliferation.

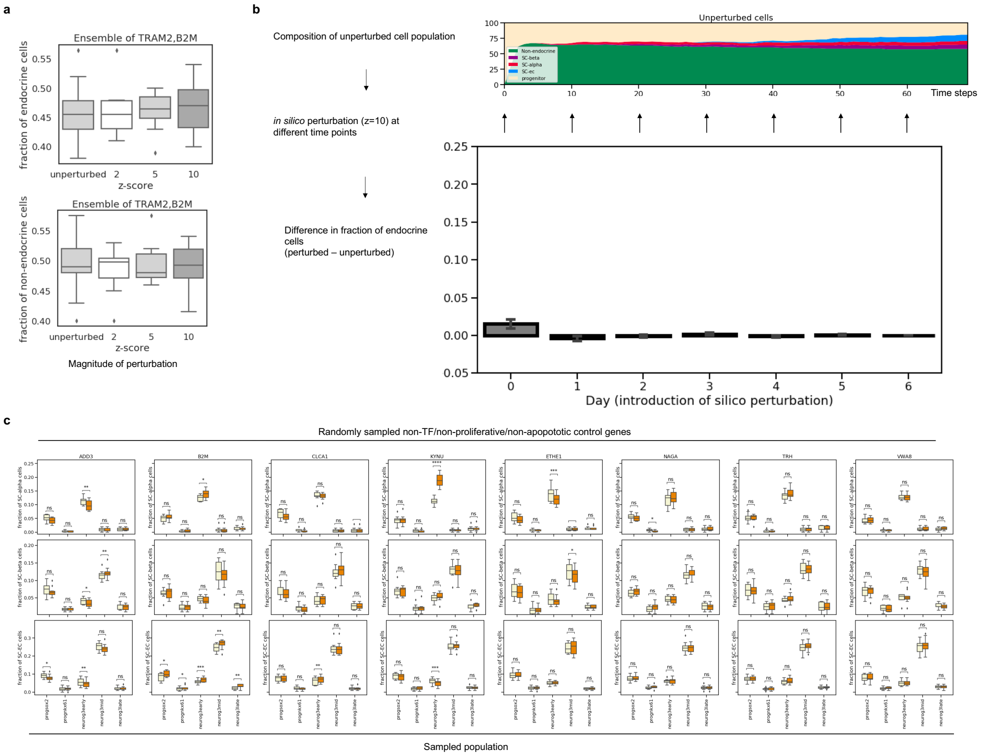


Figure S4: **PRESCIENT predicts non-significant changes to final cell fraction in response to in silico perturbations of the non-TFs not involved in apoptosis/proliferation.**

(a) Final fractions of endocrine and exocrine cells as a result of ensembled perturbations of non-TF control genes (top, bottom) on day 0. (b) in silico perturbations are introduced at different time points to the corresponding unperturbed population. The different outcomes of the cell type of interest are then calculated as the difference in fraction at the final time point starting from the perturbed and unperturbed populations. (c) Final fractions of , and EC cells when starting from different perturbed vs. unperturbed populations. Asterisks indicate significance at paired t-test p * $< 0.05$ ** $< 0.01$, *** $< 0.001$. (a-c) Results are reported over 10 randomly sampled starting populations.

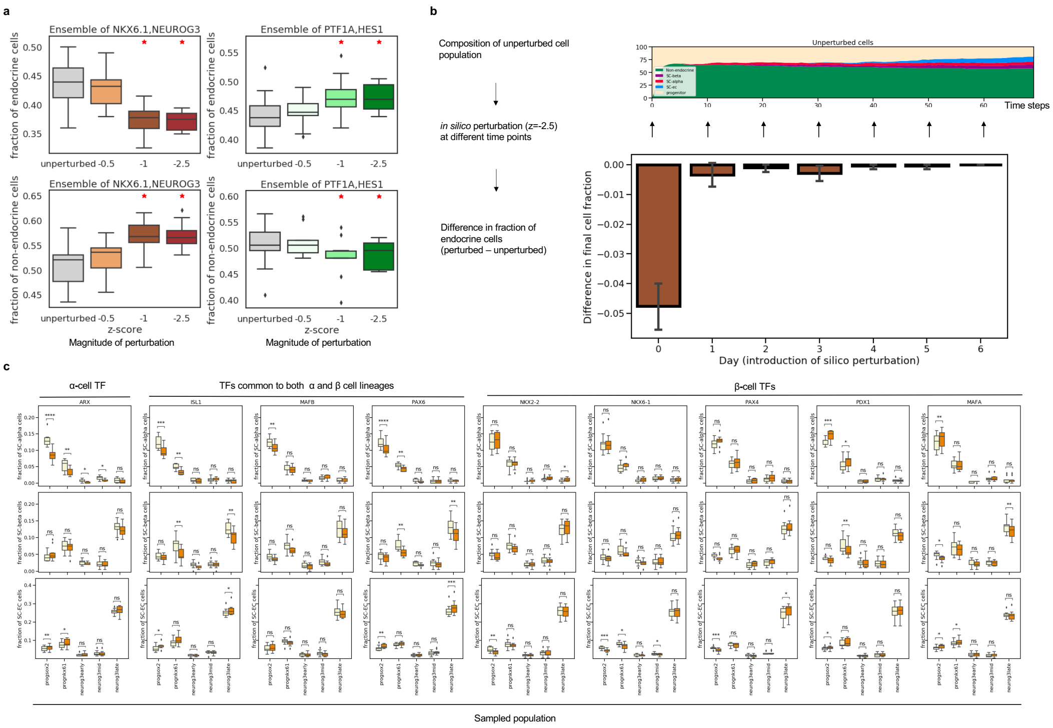


Figure S5: **PRESCIENT predicts expected outcomes of in silico knockdowns of the endocrine/exocrine axis.**
(a) Final fractions of endocrine and exocrine cells as a result of ensembled knockdowns of endocrine- and exocrine- associated TFs (left, right) on day 0 (b) in silico perturbations are introduced at different time points to the corresponding unperturbed population. The different outcomes of the cell type of interest are then calculated as the difference in fraction at the final time point starting from the perturbed and unperturbed populations. (c) Final fractions of , and EC cells when starting from different perturbed vs. unperturbed populations. Asterisks indicate significance at paired t-test p * $<$ 0.05 ** $<$ 0.01, *** $<$ 0.001. (b-c) Results are reported over 10 randomly sampled starting populations (a-c) Results are reported over 10 randomly sampled starting populations.
